# STAT3 localizes in mitochondria-associated ER membranes instead of in mitochondria

**DOI:** 10.1101/2019.12.18.880567

**Authors:** Yixun Su, Xiaomin Huang, Zhangsen Huang, Taida Huang, Yunsheng Xu, Chenju Yi

**Affiliations:** The Seventh Affiliated Hospital of Sun Yat-sen University, Shenzhen, 518107, China; Department of Biochemistry, Yong Loo Lin School of Medicine, National University of Singapore, Singapore 119615, Singapore

**Author notes:** To whom correspondence should be addressed: Prof. Chenju Yi, The Seventh Affiliated Hospital of Sun Yat-sen University, Shenzhen, 518107, China;.

**Keywords:** MAM, transcription factors, mitochondrial localization

## Abstract

Signal Transducer and Activator of Transcription 3 (STAT3) is a transcription factor (TF) that regulates a variety of biological processes, including a key role in mediating mitochondrial metabolism. It has been shown that STAT3 performs this function by translocating in minute amounts into mitochondria and interacting with mitochondrial proteins and genome. However, whether STAT3 localizes in mitochondria is still up for debate.

To decipher the role of mitochondrial STAT3 requires a detailed understanding of its cellular localization. Using Percoll density gradient centrifugation, we surprisingly found that STAT3 is not located in the mitochondrial fraction, but instead, in the mitochondria-associated endoplasmic reticulum membrane (MAM) fraction. This was confirmed by sub-diffraction image analysis of labeled mitochondria in embryonic astrocytes. Also, we find that other TFs that have been previously found to localize in mitochondria are also found instead in the MAM fraction. Our results suggest that STAT3 and other transcriptional factors are, contrary to prior studies, consolidated specifically at MAMs, and further efforts to understand mitochondrial STAT3 function must take into consideration this localization, as the associated functional consequences offer a different interpretation to the questions of STAT3 trafficking and signaling in the mitochondria.

## Introduction

Signal Transducer and Activator of Transcription 3 (STAT3) is a transcription factor (TF) encoded by the Stat3 gene in mouse. STAT3 has been found to be crucial to regulating a variety of biological processes such as embryonic development, immunogenic response, and carcinogenesis [1]. These processes occur through ligand-mediated activation of STAT3 such as through cytokines and growth factors (1). Structurally, STAT3 oligomerizes into homo- or hetero-dimers, translocating into the nucleus where it acts as a transcription activator in this form [1-3].

Over the last decade, it has been reported that STAT3 can translocate into mitochondria, where it promotes mitochondrial respiration by interacting with various mitochondrial proteins, as well as the mitochondrial genome [4-8]. Despite these findings, whether STAT3 localizes in mitochondria is still up for debate, and underlies further questions on how STAT3 can regulate mitochondrial metabolism.

In this report, we provide evidence that STAT3 does not exist in mitochondria but instead, localizes in mitochondria-associated endoplasmic reticulum membrane (MAM) in mouse brain, lung, and liver. MAM is a cellular structure formed by non-covalent protein interactions between the endoplasmic reticulum (ER) and mitochondria, which has broad implications in mitochondrial bioenergetics and reactive oxygen species production [9, 10]. This suggests that STAT3 might regulate mitochondrial metabolism via MAM function. In addition, we show that other transcriptional factors that were previously reported to be in mitochondria only exist within the MAM fraction.

## Results

### STAT3 does not localize to pure mitochondria

To determine the localization of STAT3 in mitochondria, we first sought to isolate mitochondria from primary neural progenitor cells by sucrose gradient centrifugation (Fig. 1a). STAT3 protein was detected in the mitochondria fraction, consistent with previous reports (Fig. 1b). However, the presence of ER and cytosol markers suggests contamination by the MAM fraction (Fig. 1b).

**Fig. 1.**
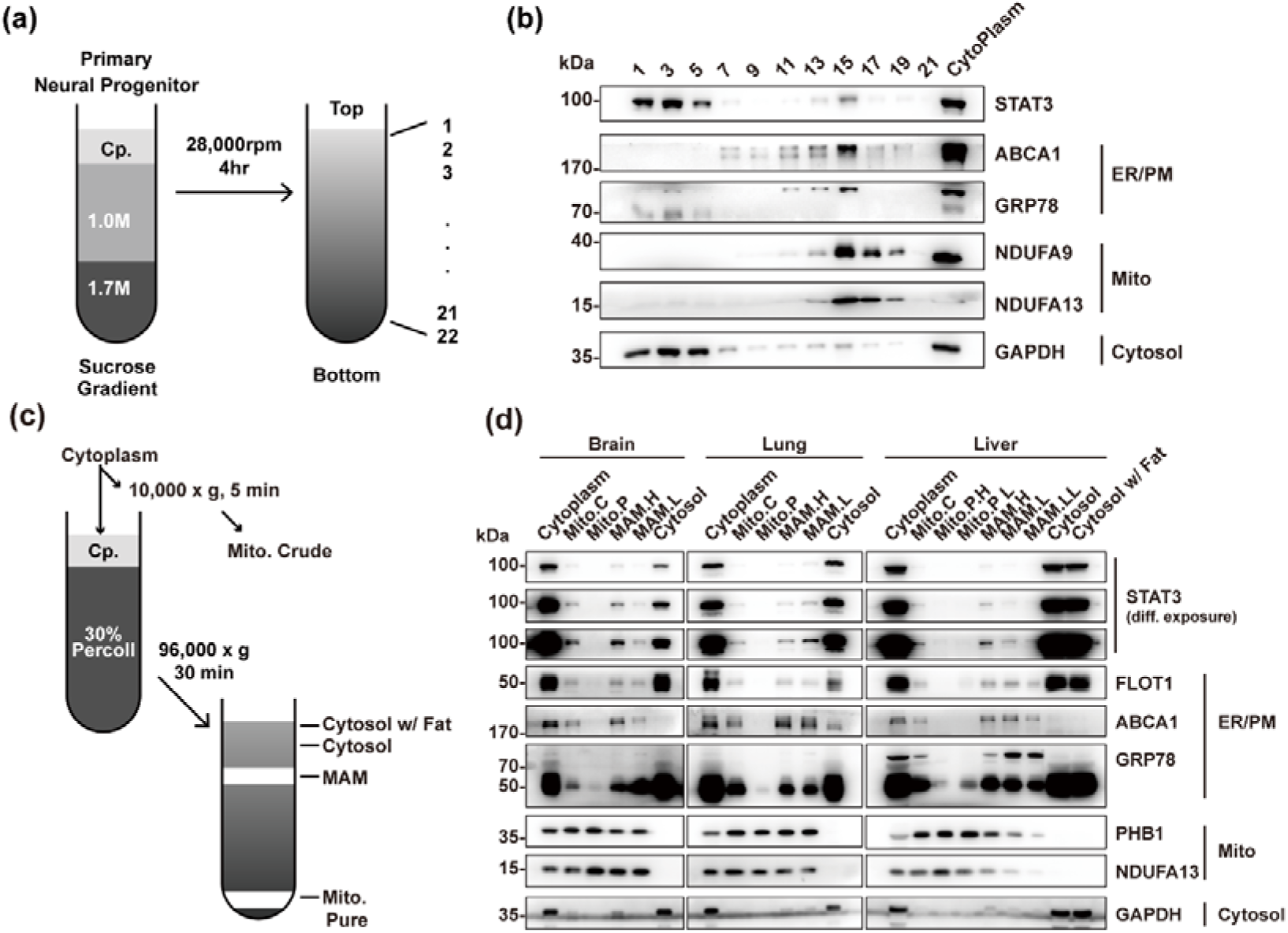
Percoll cell fractionation demonstrates that STAT3 does not exist in the mitochondria. (a) Experimental flow-chat for cell fractionation. (b) Western blots showed that a trace of STAT3 in the mitochondria fractions (15), as indicated by the existence of NDUFA9 and NDUFA13. However, ER/PM and Cytosol markers were also found in the mitochondria fraction. (c) Experimental flow-chart of Percoll density centrifugation for MAM and pure mitochondria separation. (d) Western blot showed that pure mitochondria do not contain STAT3. The purity of mitochondria was confirmed by the absence of ER/PM marker. C.p., cytoplasm. PM, plasma membrane; Mito.C, crude mitochondrial fraction; Mito.P, pure mitochondrial fraction; MAM.H, heavy MAM; MAM.L, light MAM; MAM.L.L, light-light MAM.

Percoll gradient centrifugation has been shown to be able to isolate pure mitochondria from those attached to MAM [11, 12]. Thus, we attempted to use this method to isolate pure mitochondria fractions in different mouse tissues including the brain, lung, and liver (Fig 1c). Western blot results showed that the pure mitochondrial fraction contained no detectable STAT3 protein in all three tissues, suggesting that STAT3 does not exist in mitochondria. The absence of ER and cytosol markers confirmed the successful isolation of pure mitochondria (Fig. 1d). Instead, STAT3 protein was found in the fractions containing MAM.

To further illustrate that STAT3 does not exist in mitochondria, we performed immunofluorescence studies on primary astrocytes. Using sub-diffraction image analysis (Zeiss Airyscan), we found that STAT3 did not colocalize with the mitochondrial marker heat shock protein 60 (HSP60) (Fig. 2a, b). Small amounts of STAT3 were found to localize near mitochondria, which cannot be resolved using normal confocal microscopy methods (Fig. 2b). Quantification of colocalization using the Coloc 2 colocalization analyzer found that the degree of STAT3 and HSP60 colocalization was similar to that of the negative control (DAPI and HSP60) (Fig. 2c). In contrast, STAT3 was found to colocalize with the ER (Fig. 2d∼f).

**Fig. 2.**
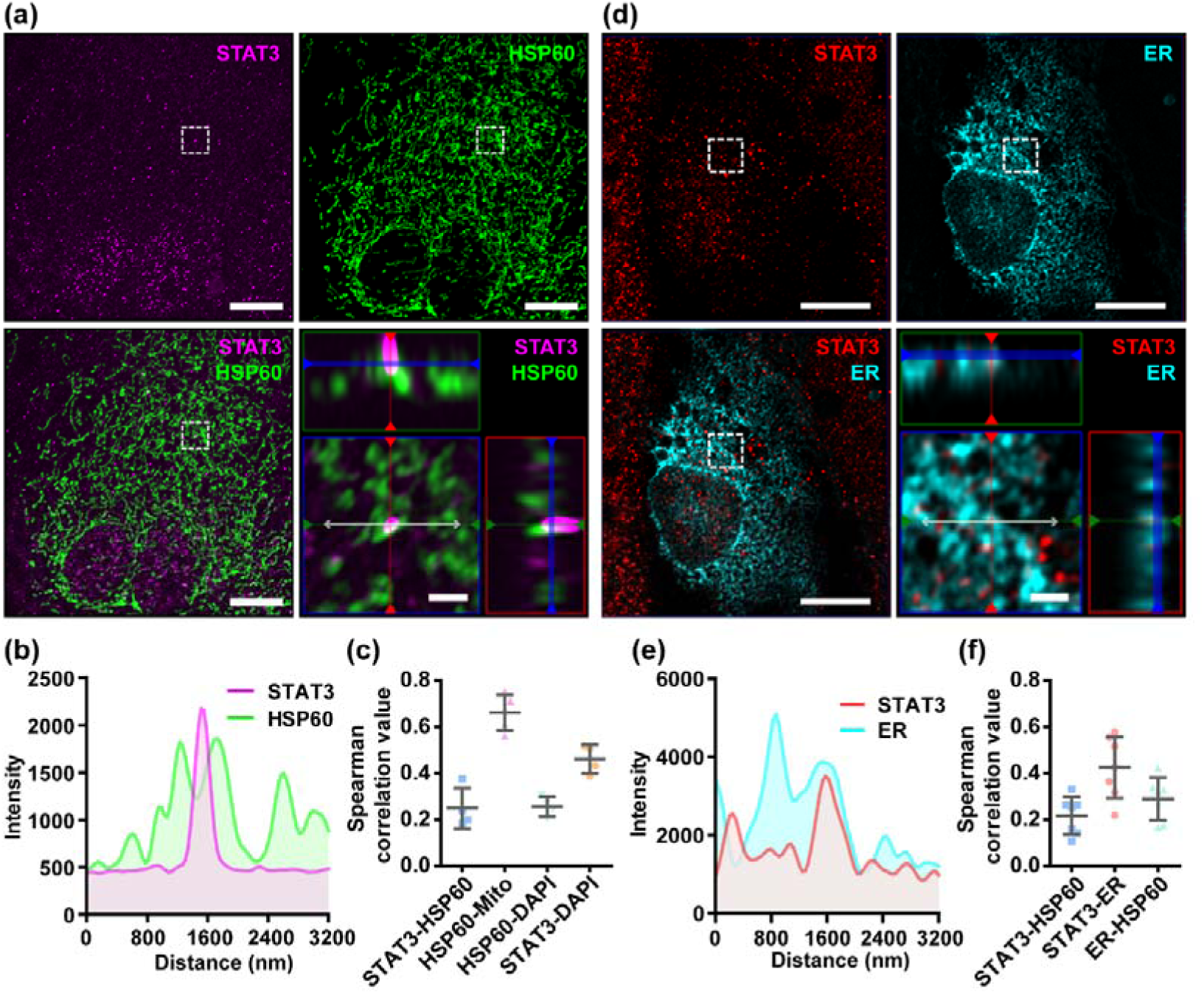
Immunofluorescence of primary astrocytes demonstrates that STAT3 does not colocalize with mitochondria. (a) Representative sub-diffraction maximum-intensity-projection image of immunofluorescence labeling of STAT3 and HSP60 in primary astrocytes (Scale bar: 10 um). The bottom-left figure shows the orthogonal views of the highlighted areas (Scale bar: 1 um). (b) The intensity of STAT3 and HSP60 on the double-headed line in (a). (c) Correlation coefficients between STAT3, HSP60, MitoTracker DeepRed (Mito) and DAPI. (d) Representative sub-diffraction maximum-intensity-projection image of immunofluorescence labeling of STAT3 and ER-dsRed (ER) in primary astrocytes (Scale bar: 10 um). The bottom-left figure shows the orthogonal views of the highlighted areas (Scale bar: 1 um). (e) The intensity of STAT3 and ER-dsRed on the double-headed line in (d). (f) The correlation coefficient between STAT3, HSP60, and ER-dsRed.

In addition, the number of STAT3 puncta observed per cell was found to be 170±14, which corroborates previous reports (the number of STAT3 molecules per cell is 103∼146 via Western Blotting, 27∼126 via mass spectrometry) [13], verifying the reliability of our immunostaining results.

### Other methods to examine STAT3 localization in mitochondria were not able to completely remove MAM

In prior work done by other groups, several pure mitochondria isolation methods have been used to evaluate the localization of STAT3 in mitochondria, including trypsinization and sonication. We sought to examine whether these methods can reliably dissociate the MAM fractions from the pure mitochondria fractions, and analyzed whether STAT3 localizes with ER/cytosol markers in these fractions.

Sonication methods resulted in the disruption of not only MAM but also mitochondria, as shown by a decreased level of both markers in the pellet (Fig. 3a). In contrast, trypsinization had little deleterious effects on mitochondria integrity, but only achieved partial removal of MAM despite long incubations with enzyme of up to 60 min (Fig. 3b). In separate studies, it has been reported that high salt washes of mitochondria followed by trypsinization can disrupt protein interactions and dissociate attached actin filaments [14]. We attempted to examine if this method was able to remove the attached MAM from mitochondria. The results demonstrated that high salt washes combined with trypsinization were still unable to obtain pure mitochondria (Fig. 3c). Though STAT3 remained in the mitochondria fractions obtained by all these methods, this may be explained by the presence of MAM fraction remnants, as indicated by the contamination of MAM and cytosol markers (Fig. 3a∼c).

**Fig. 3.**
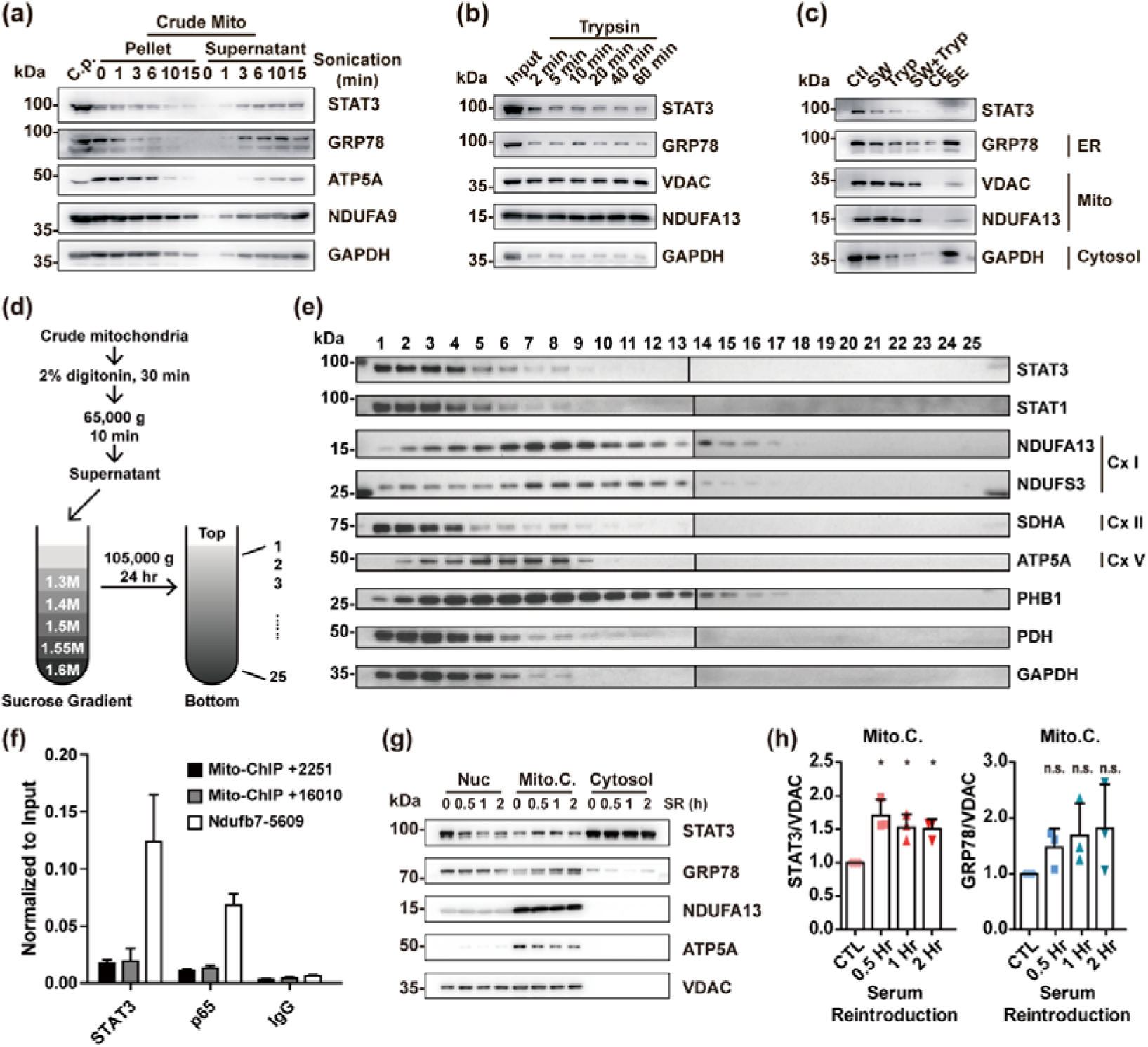
Re-examining the existence and function of ‘mitochondrial STAT3’. (a) Purification of mitochondria by sonication. (b) Purification of mitochondria by trypsinization. (c) Purification of mitochondria by washing with high concentration of salt combined with trypsinization. GRP78 was used as the ER marker; ATP5A, NDUFA9, NDUFA13, and VDAC were used as the mitochondrial marker; GAPDH was used as the cytosolic marker. (d) Experimental flow-chart of mitochondrial fractionation. (e) Western blot analysis of mitochondrial fractions. (f) ChIP-qPCR detection of STAT3-binding on mitochondrial DNA in mouse embryonic stem cells. (g) Serum reintroduction experiment in Neuro2A cells. (h) Quantification of Western blot results in (g) in triplicates. C.p., cytoplasmic fraction. Ctl, control; SW, salt-washed; Tryp, Trypsinized; CE, control elute; salt-washed elute; Cx I, complex I; Cx II, complex II; Cx V, complex V; Nuc, nuclear fraction; Mito.C., crude mitochondrial fraction; SR, serum reintroduction.

In conclusion, the aforementioned methods fail to perfectly isolate pure mitochondria, and thus are unable to confirm the exclusive localization of STAT3 to mitochondria. At the same time, these results demonstrate that the Mitochondria-ER contacts may be resistant to sonication, trypsinization, and high salt washing.

### STAT3 does not colocalize with Complex I, and its level correlates with the MAM level

While we have demonstrated that STAT3 does not exist within mitochondria, other studies have shown that mitochondrial STAT3 binds to Complex I and to mitochondrial DNA to modify mitochondrial metabolism [6, 8]. However, our fractionation analysis demonstrates that STAT3 does not exist within the Complex I fraction (lane 7, 8; Fig. 3d∼e). In addition, our ChIP-qPCR experiments demonstrate that STAT3 does not bind to mitochondrial DNA in mouse embryonic stem cells (Fig. 3f).

Previous studies have found that STAT3 levels in crude mitochondria increase after serum reintroduction following serum starvation [4]. We hypothesized that this may result from increased MAM levels [15]. We performed serum reintroduction studies on Neuro2A cells. We found that reintroduction of serum increased the amount of STAT3 in the mitochondria fraction at as early as 30 min (Fig. 3g, h), accompanied by an increased level of ER marker glucose-regulated protein 78 (GRP78) in the same fraction (Fig. 3g, h), suggesting that increased STAT3 levels may be due to the increase of MAM in the crude mitochondrial fraction.

### Other TFs or signaling proteins are present in MAM, but not in mitochondria

In addition to STAT3, a number of other TFs or signaling proteins have been previously found to localize in mitochondria where they are involved in regulating mitochondrial function, including STAT1 [16, 17], mitogen-activated protein kinase 1/3 (MAPK1/3) [18, 19], MAPK14 [20, 21], AMP-activated protein kinase (AMPK) [22], protein kinase B (AKT) [23, 24], mammalian target of rapamycin (mTOR) [24], as well as NF-κB p65 subunit (RELA) [25, 26]. We wondered whether, like STAT3, they actually localize in MAM instead of within mitochondria. We examined their localization in the pure mitochondrial fraction using Western blotting. The results demonstrate that none of these TFs were detectable in the pure mitochondria fraction (Fig. 4). Catenin beta 1(CTNNB1), which regulates mitochondrial metabolism but has never been found in mitochondria [27], was also found in the MAM fraction, suggesting a possible mechanism for MAM to regulate mitochondrial metabolism (Fig. 4).

**Fig. 4.**
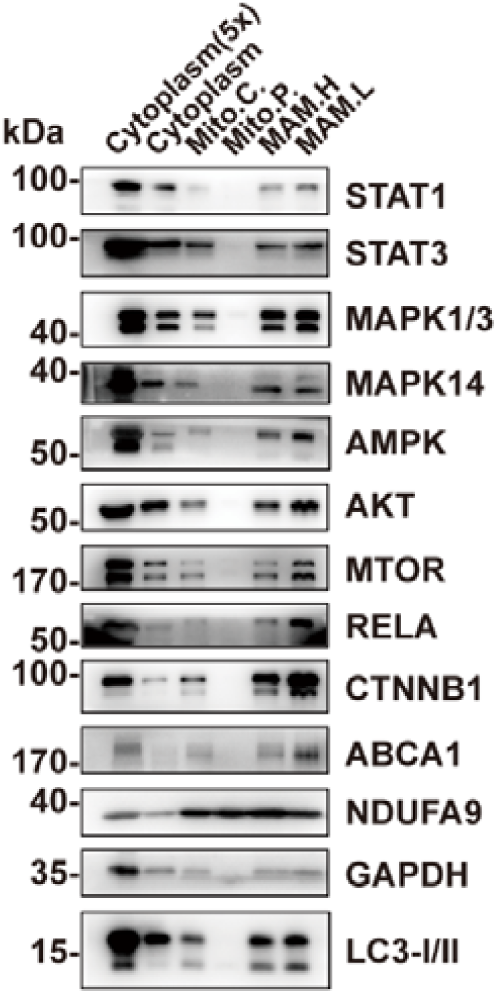
Other TFs or signaling proteins are present in MAM but not in mitochondria. Western blot analysis of the Percoll density centrifugation result showed that TFs or signaling proteins including STAT1, MAPKs, AMPK, AKT, mTOR, and RELA could only be found in the MAM fractions but not in pure mitochondria. Mito.C., crude mitochondrial fraction; Mito.P., pure mitochondrial fraction; MAM.H, heavy MAM; MAM.L, light MAM.

## Discussion

STAT3 has been shown to regulate mitochondrial metabolism by translocating to mitochondria [6, 8]. Initially, our study intended to investigate the functional mechanism of STAT3 in mitochondria. However, our work demonstrates that STAT3, along with some other nuclear TFs, are not present in the mitochondria, but instead are found in the MAM. This suggests that STAT3 may regulate mitochondrial function via other mechanisms yet unknown. It has been reported that STAT3 can be found in the endoplasmic reticulum (ER) compartment, where it interacts with the inositol trisphosphate receptor (IP3R) and controls calcium efflux from the ER [28]. IR3R has also been found to form the MAM contact site by interacting with voltage-dependent anion channels (VDAC) on the mitochondrial outer membrane [29]. Thus, it is possible that MAM-STAT3 may regulate mitochondrial metabolism through modulating calcium transport. Another possibility is that TFs and other nuclear proteins are only transported to the MAM fraction for degradation [30].

Several questions surrounding the “mitochondria-STAT3” theory have remained unanswered in the past decade. Firstly, what is the mechanism of the mitochondrial-import of STAT3 and other nuclear TFs? Protein import into mitochondria is restricted by the impermeable double membrane of mitochondria, and only proteins with mitochondrial signaling peptides can be imported into mitochondria through complex molecular machinery during translation [31, 32]. However, little evidence exists that uncovers the mechanism of STAT3 import. It has been suggested that NADH: ubiquinone oxidoreductase subunit A13 (NDUFA13) is required for STAT3 import to mitochondria [33], yet the key mitochondrial channel responsible for this action has not been identified [34]. Until further evidence is uncovered, mitochondrial import of STAT3 is not well-supported by experimental results.

Secondly, how does “mitochondria-STAT3” regulate mitochondrial metabolism? Several mechanisms have been proposed, including its interaction with respiratory complex I/II, cyclophilin D (CypD) and pyruvate dehydrogenase, as well as its regulation of the mitochondrial transcription [4-8]. However, even if STAT3 was localized in mitochondria, it is unlikely to influence mitochondrial metabolism through direct interaction with mitochondrial proteins or genome, considering the stoichiometric difference between STAT3 (∼10^2^ molecules/cell) and mitochondrial protein (6 x 10^6^ molecules of Complex I or II/cell) or mitochondrial genome (1000∼5000 copies/cell) [13, 35, 36]. Proposed mechanisms of STAT3 activity by other groups are largely based on the overexpression strategies of “mito-STAT3” (STAT3 fused with a mitochondrial signaling peptide at the N-terminus) which employs the assumption that STAT3 localizes to mitochondria [4, 5, 8]. These results may prove to be artificial or artifactual if this assumption proves to be false.

In summary, our results suggest that STAT3, along with other TFs or signaling proteins with previously described mitochondrial localization, actually localize in MAM, contrary to prior reports. Extra attention must be paid to the localization of these factors in the future due to the inherent subjectivity of cell fractionation.

## Materials and Methods

### Animal

Wild type C57BL/6j mice were euthanized by CO_2_ overdose before dissection and tissues or embryos collection. All animals used in the study were maintained in the Specific-pathogen-free animal facility. All procedures were performed under the approval of the Sun Yat-sen University Institutional Animal Care and Use Committee.

### Cell culture

Primary neural progenitors were isolated from embryonic day (E) 14 mouse embryos and cultured in serum-free medium (Dulbecco’s Modified Eagle Medium/ Nutrient Mixture F-12 (DMEM/F12) with B27 supplement and epidermal growth factor (EGF)/ fibroblast growth factor (FGF). Astrocytes were isolated from E15 embryos and were cultured in DMEM with 10% FBS. Neuro2A cells were cultured and passaged in DMEM with 10% FBS. Serum reintroduction experiments were performed by culturing cells in serum-free DMEM media for 6 hours, followed by serum reintroduction at various time points prior to collection.

### Cell fractionation

Cells cultured in 10 cm dishes were scraped down in 0.5 ml isotonic buffer (5 mM HEPES, 250 mM sucrose, 0.1 mM EDTA) and transferred to an Eppendorf tube. The dishes were then washed with 0.5 ml isotonic buffer, which was then pooled together. Cells were homogenized by repeated passes through a 25-gauge needle attached to a 1-ml syringe 15 times. 100 µl of the homogenate was saved as the whole cell lysate control. The remaining homogenate was centrifuged at 750 g for 5 min to remove intact cells and nucleus. The cytoplasm was collected from the supernatant, and 100 µl was saved as control. The cytoplasmic fraction was then layered on a sucrose gradient (7 ml 1.0 M sucrose, 3 ml 1.5 M sucrose, each contained 5 mM HEPES and 0.1 mM EDTA) in a 14 ml ultracentrifugation tube. The tubes were then subjected to ultracentrifugation in a SW40Ti rotor at 28,000 rpm for 4 h. The resulting gradient was fractionated and subjected to Western blot analysis.

### Isolation of pure mitochondria and MAM fractions

The isolation of pure mitochondria and MAM fractions from mouse tissue was performed as previously described [11, 12]. Briefly, tissues (brain, liver, muscle) were dissected from euthanized mice, followed by homogenization. Homogenates were then centrifuged at 750 g for 5 min to remove unbroken cells and nucleus. An aliquot of the cytoplasmic fraction collected from the supernatant was then centrifuged at 10,000 g for 5 min to obtain the crude mitochondria fraction. The rest of the cytoplasmic fraction was layered on 10 ml 30% Percoll buffer and centrifuged at 96,000 g for 30 min. The resulting gradient was confirmed to contain two yellow/white bands. The top band (MAM fraction) and the bottom band (pure mitochondria) were separately collected. Excess Percoll was removed from both mitochondrial and MAM by diluting the fractions with mitochondria resuspension buffer (MRB) and centrifugation at 6,000 g for 15 min, followed by supernatant centrifugation at 100,000 g for 60 min to recover light fractions. The protein concentration was measured by the Bradford assay. For Western blot analyses, 5 times of the cytoplasmic fraction (by mass) to mitochondria were loaded.

### Purification of mitochondria by sonication, trypsinization, and washing with high concentration of salt

The crude mitochondria were obtained from mouse brain as described above. To purify mitochondria by sonication, the crude mitochondria were subjected to sonication by the Scientz08-II sonicator at 50% power for 0 min, 1 min, 3 min, 6 min, 10 min, and 15 min, respectively. The resulting suspensions were centrifuged at 10,000 g for 5 min. The pellet and supernatant were collected separately. The pellets were washed once and resuspended in MRB.

To purify mitochondria by trypsinization, 2 µg/ml trypsin was added to crude mitochondrial fractions. The trypsinization was stopped at 2 min, 5 min, 10 min, 20 min, 40 min, and 60 min, respectively by FBS addition. The resulting suspensions were centrifuged at 10,000 g for 5 min and the pellets were collected and washed once and resuspended in MRB.

To purify mitochondria by washing with high concentration of salt, the crude mitochondrial fraction was incubated in MRB supplemented with 1 M KCl and 2 mM MgCl_2_ for 15 min on ice, and centrifuged at 10,000 g for 5 min. The salt-washed mitochondria were recovered in the pellet, while the eluted protein was recovered in the supernatant. Subsequently, salt-washed mitochondria were incubated in 2% trypsin for 30 min. Mitochondria were then collected by centrifugation, washed once and resuspended in MRB.

### Mitochondria fractionation

The mitochondrial fractionation protocol was adopted from Shizawa-Itoh et al. [37]. The crude mitochondria were isolated and incubated with 2% digitonin (detergent: protein = 4:1) on ice for 30 min. The sample was then centrifuged at 65,000 g for 10 min to remove the insoluble fractions. The supernatant was then loaded onto a sucrose gradient (1.3 M, 1.4 M, 1.5 M, 1.55 M, 1.6 M), and then centrifuged at 105,000 g for 24 h. The resulting gradient was then fractionated from top to bottom.

### Western Blotting

Protein samples were lysed in 5 x SDS sample buffer containing beta-mercaptoethanol, boiled at 95°C for 10min and loaded into each well for SDS-PAGE. Samples were then transferred to PVDF membrane (Thermo) and immunoblotted using anti-NDUFA9 (Invitrogen), anti-NDUFS3 (Invitrogen), anti-NDUFA13 (Invitrogen), anti-ATP5A (Invitrogen), anti-SDHA (CST), anti-VDAC (CST), anti-HSP60 (CST), anti-PHB1 (CST), anti-PDH (CST), anti-GAPDH (Sigma), anti-GRP78 (SCBT), anti-ABCA1 (SCBT), anti-FLOT1 (CST), anti-STAT3 (CST), anti-STAT1 (SCBT), anti-AMPK (CST), anti-ERK1/2 (CST), anti-p38 (CST), anti-LC3 (CST), all diluted in 5% BSA:TBST at 1:1000, followed by appropriate HRP-conjugated secondary antibodies (Thermo) incubation (diluted in 5% non-fat milk:TBST at 1:10,000), and developed using the SuperSignal™ West Femto Maximum Sensitivity Substrate (Thermo).

### Immunofluorescence

Astrocytes isolated from E15 and cultured on poly-D-lysine-coated coverslips were stained with 5 µM MitoTracker DeepRed to label mitochondria. Alternatively, cells were transfected with ER-dsRed expression vector one day prior to fixation to label ER. Cells were fixed with 4% paraformaldehyde for 15min and were then permeabilized by ice-cold methanol for 5 min, followed by PBS rinse for 10 min. The cells were then blocked with 5% bovine serum albumin in PBS with 0.3% TritonX-100 for 1 hr, followed by primary antibody (rabbit-anti-HSP60 (CST), mouse-anti-STAT3 (CST), both diluted at 1:200 in the blocking buffer) incubation at 4°C overnight. Then, the cells were incubated in appropriate secondary antibodies (goat-anti-rabbit-AF488, goat-anti-mouse-AF568 or goat-anti-mouse-AF647 (Abcam), both diluted at 1:200 in the blocking buffer) for 1h at room temperature. After washing 3 times in TBST, the cells on the coverslip was mounted using Prolong Gold with DAPI (Thermo) and subjected to confocal microscopy observation using the Zeiss LSM880 with Airyscan system. Images were captured using a 63x/1.4NA oil immersion objective. ER-dsRed was detected using 543 nm laser. HSP60 was detected using 488 nm laser. STAT3 was detected using 543nm or 631 nm laser (depending on the secondary antibody used). MitoTracker DeepRed was detected using 631 nm laser. The colocalization analysis was performed with Fiji software using the Coloc 2 plugin.

### ChIP-qPCR

Mouse embryonic stem cells were crosslinked with formaldehyde at a final concentration of 1% for 10min followed by quenching with glycine. Chromatin extracts were fragmented by sonication and pre-cleared with protein G Dynabeads, then subsequently precipitated with anti-STAT3 antibody (Santa Cruz), anti-p65 antibody (Santa Cruz), or normal rabbit IgG (Santa Cruz) overnight at 4°C. After washing and elution, crosslink reversal was done by incubating at 65°C for 8h. The eluted DNA was purified and analyzed by qPCR with primers specific to the predicted STAT3 binding site. qPCR experiment was carried out using SYBR Green qPCR kit from KAPA and Applied Biosystems 7500 Real PCR System. Samples were assayed in duplicate. Primers sequences are: Mt-ChIP-2F, tggggtgacctcggagaat; Mt-ChIP-2R, cctagggtaacttggtccgt; Mt-ChIP-13F, ccgcaaaacccaatcacctaag; Mt-ChIP-13R, ttggggtttggcattaagagga; mNdufb7-F, tctgttaaatgtcacccgtcct; mNdufb7-R, acttttacacctggtacccaaca.

### Statistic

Statistical significance was determined by Student’s t-test using GraphPad Prism 6.01. The p value < 0.05 was considered significant. Unless otherwise specified, data were presented as mean and the standard deviation (mean ± SD).

## Acknowledgments

This work was supported by grants from the National Nature Science Foundation of China (NSFC 81971309) and Guangdong Province Basic Research, Applied Basic Research Foundation (2019A1515011333) and Sun Yat-sen University Key Training Program for Youth Teachers (F7201931620002). We would like to thank Dr. Jason Tann and Prof. Hui Chen for revising the manuscript.

## Author contributions

**Yixun Su:** Conceptualization, Methodology, Visualization, Investigation, Writing - Original Draft. **Xiaomin Huang:** Investigation. **Zhangsen Huang:** Investigation. **Taida Huang:** Validation. **Yunsheng Xu:** Supervision. **Chenju Yi:** Supervision, Writing – Review & Editing, Funding acquisition.

## Abbreviations

STAT3: Signal Transducer and Activator of Transcription 3
MAM: mitochondria-associated endoplasmic reticulum membrane
TF: transcription factor
ER: endoplasmic reticulum
HSP60: heat shock protein 60
GRP78: glucose-regulated protein 78
MAPK: mitogen-activated protein kinase
AMPK: AMP-activated protein kinase
AKT: protein kinase B
mTOR: mammalian target of rapamycin
RELA: NF-κB p65 subunit
CTNNB1: Catenin beta 1
IP3R: inositol trisphosphate receptor
VDAC: voltage-dependent anion channels
NDUFA13: NADH: ubiquinone oxidoreductase subunit A13
CypD: cyclophilin D

